# Plasticity of squamous differentiation drives drug resistance in HNSCC

**DOI:** 10.64898/2026.03.09.710514

**Authors:** Kalle Sipilä, Matteo Vietri Rudan, Priyanka Bhosale, Matthew Blakeley, Clarisse Ganier, Robert Kennedy, Emanuel Rognoni, Fiona M. Watt

## Abstract

A critical hallmark of carcinogenesis is the ability of cancer cells to evade the loss of self-renewal normally imposed by terminal differentiation. However, therapies directly attempting to promote differentiation have shown limited efficacy in solid tumours and the cellular mechanisms behind cancer cell persistence are poorly understood. Here we established a patient-derived orthotopic head and neck squamous cell carcinoma (HNSCC) model in vivo that recapitulates the genetic, cellular and histopathological heterogeneity of HNSCC. Experimental induction of differentiation and clonal lineage tracing by fluorescent barcoding revealed a heterogeneous response to terminal differentiation stimuli, enabling subsets of cancer cells to escape differentiation-associated loss of self-renewal. While pharmacological inhibition of ErbB-MEK1/2-ERK1/2 pathway by afatinib could induce the differentiation of patient-derived cancer cells, some highly clonogenic cells remained refractory to differentiation signals even though they were capable of differentiating. Differentiation reporter IVLmCherry further confirmed that differentiation and loss of self-renewal ability were partially uncoupled in patient-derived HNSCC cells. These findings identify differentiation-resistant clonogenic populations as a key barrier to therapeutic efficacy and provide a framework for improving differentiation-based strategies in HNSCC.

## Introduction

Head and neck squamous cell carcinoma (HNSCC) is the seventh most common cancer worldwide, and its five-year survival remains poor (Creaney et al., 2022). HNSCC is frequently associated with human papillomavirus (HPV) infection, whereas the principal risk factors for HPV-negative (HPV^-^) disease are smoking and alcohol consumption. HPV-positive HNSCC is typically associated with a more favourable prognosis compared with HPV^–^ disease. Surgical resection combined with chemoradiotherapy remains the standard treatment strategy for most patients *with HPV - HNSCC*. Few targeted therapies have demonstrated clinical efficacy in HNSCC. Cetuximab, an epidermal growth factor receptor (EGFR)-blocking antibody, can be administered in combination with chemoradiotherapy (Johnson et al., 2020). Interestingly, PD-1 targeting immunotherapy has recently been shown to improve overall survival (Uppaluri et al., 2025).

It is widely recognised that a key component of carcinogenesis is the ability of cancer cells to evade the loss of self-renewal associated with terminal differentiation (Hanahan, 2022). In normal stratified squamous epithelia, proliferative cells reside in the basal layer, where they are attached to the basement membrane. Upon commitment to differentiation, cells move to the suprabasal layers and initiate expression of differentiation markers, including cornified envelope precursor proteins such as involucrin (IVL) and loricrin. During this process, squamous epithelial cells withdraw from the cell cycle and ultimately lose their self-renewal capacity although proliferation and differentiation are controlled separately (Watt, 1988). HNSCC is genetically highly heterogeneous and shows frequent mutations in the squamous differentiation pathway, including loss-of-function mutations of Notch1, P53, Casp8, and PIK3CA (Cancer Genome Atlas Network 2015). HPV^-^ HNSCC are usually either well or moderately differentiating while well differentiated tumours show better prognosis (Johnson et al., 2020). Interestingly, recent single cell and spatial analysis have identified new phenotypic features of stromal composition and partial epithelial-mesenchymal transition that are associated with clinical outcomes (Punovuori at al., 2024).

Inducing terminal differentiation to drive irreversible loss of self-renewal represents a potentially attractive therapeutic strategy proven to be effective treatment in some cancers, such as acute promyelocytic leukaemia (APL) (de Thé, 2018). In APL, the PML–RARα fusion protein blocks granulocytic differentiation. Treatment with all-trans retinoic acid (ATRA) restores granulocytic differentiation and leads to the clearance of the majority of leukemic cells, leading to rapid but often transient remission. Importantly, permanent cure of APL requires high-dose ATRA or arsenic trioxide, which induces sustained loss of self-renewal of tumour maintaining cells through destabilisation and degradation of the fusion protein (de Thé, 2018). These observations highlight the importance of understanding how distinct cellular subpopulations within a heterogeneous tumour respond to differentiation cues. In addition, epithelial cells display some naturally-occurring plasticity that can interfere with the development of differentiation therapies for keratinocyte cancers. Wounding studies in normal epithelium have demonstrated that Gata6+ differentiated cell population in sebaceous ducts can retain the capacity to de-differentiate and reacquire self-renewal potential (Donati et al., 2017). It is unclear whether this is relevant to the head and neck mucosa.

The commitment to differentiation in keratinocytes is regulated by a complex protein phosphatase network targeting ErbB, MAPK, insulin, and adhesion signalling (Mishra et al., 2017; Hiratsuka et al., 2020). Interestingly, alterations of these pathways have been shown to be common feature in HNSCC tumours (Punovuori et al., 2024; Puram et al., 2017; Rheinwald and Beckett, 1980); furthermore HNSCC cells in culture can be induced to undergo differentiation by EGFR inhibitors (Setúbal Destro Rodrigues et al., 2018). However, understanding the complex changes in cell behaviour in vivo requires characterization of differentiation dynamics, stemness, and self-renewal within the heterogeneous cancer cell population in cancer models in vivo. Here, we apply epithelial stem cell biology methods to establish a patient-derived in vitro and in vivo model of HNSCC that captures the mutational and cellular heterogeneity of HNSCC cancer cells. We demonstrate that while experimental induction of differentiation by methylcellulose suspension assay or by blocking ErbB with afatinib induces some cancer cells to undergo differentiation, the clonogenic tumour initiating cells show a striking resistance to differentiation signals and do not lose the capability to form differentiative progeny in lineage tracing experiments. This suggests that differentiation plasticity is an important intrinsic mechanism of drug resistance that must be overcome for HNSCC differentiation therapies to be effective in the clinic.

## Results

### Patient-derived cancer cells recapitulate the histopathological complexity of HPV^-^ HNSCC in xenografts

We previously created a library of mutationally heterogeneous cancer cells from HPV^-^ HNSCCs (SJG lines; each SJG line was established from a single patient), which are maintained in vitro by using J2-3T3 feeder cells in a system similar to epidermal stem cell culture (Hayes et al., 2016). This feeder-based approach enables robust expansion of cancer cells while reducing culture-driven clonal selection. Analysis of the somatic mutational landscape, defined relative to matched blood controls, demonstrated that these cells recapitulate the heterogeneous genomic features typical of HNSCC, including frequent mutations in TP53, PIK3CA, FAT1, NOTCH1 and CDKN2A (Hayes et al., 2016, Fig. 1B). Immunofluorescence staining for the basal epithelial marker keratin 14 (K14) and the differentiation marker involucrin (IVL) revealed that different SJG lines (i.e. from different patients) show substantial variation in the expression levels of IVL as well as in the way how they assemble the basal and differentiating cell compartments in vitro (Fig. 1C).

**Figure 1.**
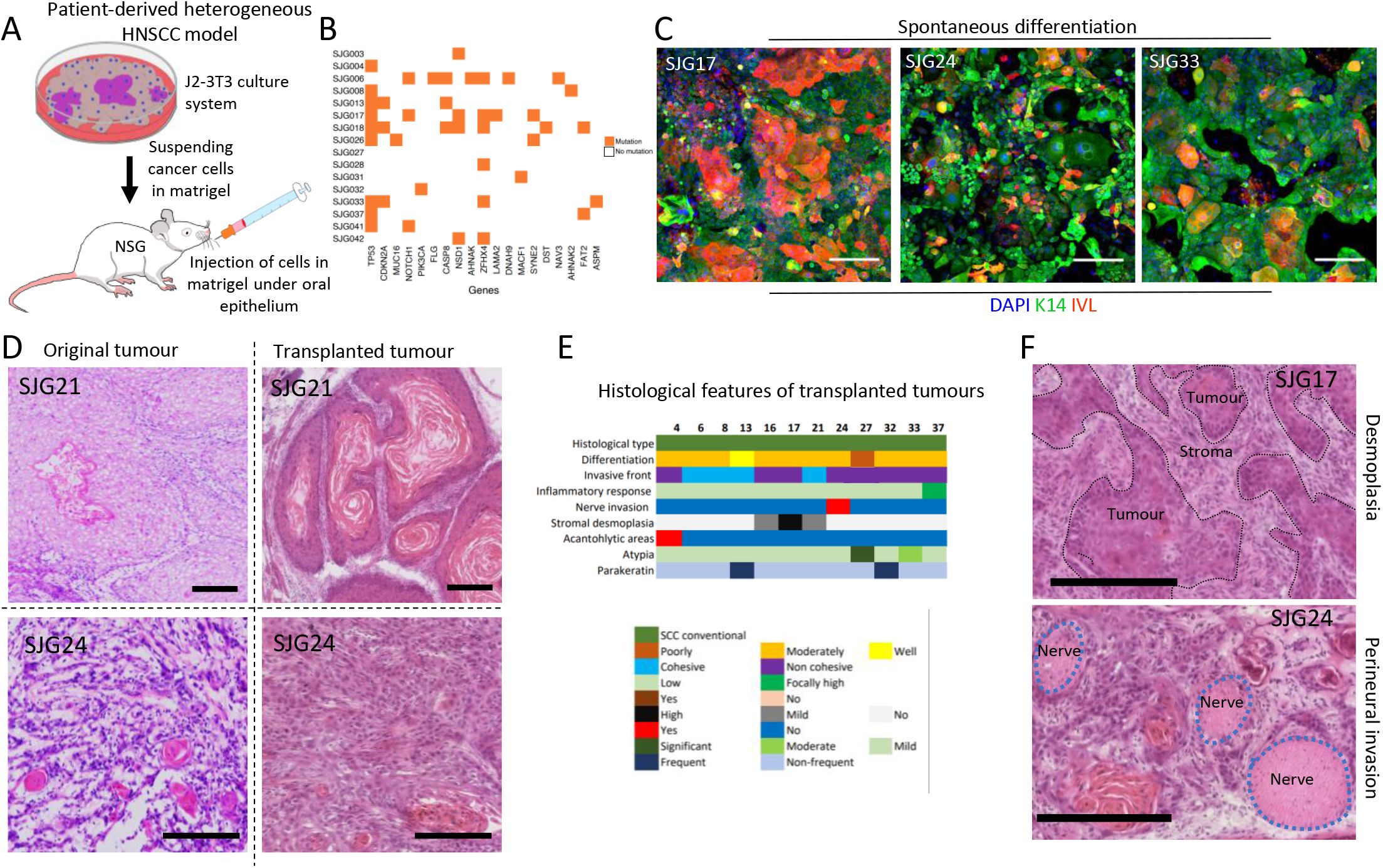
An orthotopic patient-derived HPV-negative HNSCC model recapitulates the pathological and cellular heterogeneity of the disease. (A)Schematic overview of the patient-derived HNSCC model. (B)Somatic exon mutations identified in different SJG lines (Redrawn from Hayes et al., 2016). (C)SJG13, SJG24, or SJG33 cells co-cultured with J2-3T3 cells and stained for K14 (green), IVL (red), and nuclei (DAPI, blue). Note variation in basal cell morphology and proportion of IVL expressing cells. Scale bars: 250μm. (D)Haematoxylin and eosin (H&E) staining of transplanted SJG21 and SJG24 tumour cryosections alongside their corresponding original tumours. Scale bars: 250μm (E)Summary of histopathological features observed in the library (12 different SJGs, one tumour per SJG) transplanted SJG tumours. (F)H&E staining of SJG13 and SJG24 tumours illustrating stromal desmoplasia (SJG13) and perineural invasion (SJG24).

To explore the differentiation features in vivo, we established a cancer model in which SJGs were resuspended in Matrigel and orthotopically transplanted into the cheek subepithelial compartment of immunocompromised NSG mice (NOD scid gamma mouse; Fig. 1A). 12 out of 14 tested SJGs formed rapidly growing tumours. The resulting xenografts recapitulated key histological features characteristic of human head and neck squamous cell carcinoma (HNSCC) (Fig. 1D-1F). Comparison with matched primary tumour material, where available, showed that the xenografts closely resembled the corresponding primary tumours in terms of histological architecture, organization of differentiated cell layers and stromal components, as illustrated by haematoxylin and eosin staining (H&E) of SJG21 and SJG24 primary tumours and xenografts (Fig. 1D).

Histopathological analysis showed that all SJGs formed squamous cell carcinomas but differed based on their invasive fronts, differentiation status, parakeratosis, and inflammatory response (Fig. 1E). In the case of SJG16, SJG17, and SJG21 high level of stromal desmoplasia was observed. Interestingly, in SJG24 xenografts perineural invasion was evident (Fig. 1F). Together, these findings demonstrate that the patient-derived SJG library captures the key histopathological features of HNSCC in vivo while recapitulating the significant heterogeneity among individual patients.

### Differentiation capacity of HNSCC cells in vitro does not correlate with tumour growth or clonal architecture in vivo

In normal squamous epithelium, proliferating cells are confined to the basal layer adjacent to the stroma, with differentiating cells positioned suprabasally. Cancer cells are known to show reduced dependence on the stromal niche and partial de-coupling of proliferation from commitment to differentiation (Parkinson, 1985). To study this, we utilized a methylcellulose suspension model that drives keratinocytes to undergo differentiation by preventing cell– matrix interaction (Adams and Watt, 1989).

We selected three independent patient-derived cell lines, SJG13, SJG24, and SJG33, that formed rapidly growing, moderately differentiating squamous cell carcinomas in the in vivo model, closely reflecting the histology of aggressive human HNSCC. When cultured in methylcellulose as single-cell suspensions, tumour cells exhibited increased expression of differentiation markers IVL and transglutaminase 1 (TGM1), measured by qPCR (Fig. 2A and 2B). While we did not detect an increase of IVL expression after 6h, except in the case of SJG13, IVL expression was significantly increased after 20h in normal oral keratinocytes (OK) as well as SJG13, SJG24, and SJG33 (Fig. 2A). In contrast, TGM1 expression increased at 6h in OK and SJG lines (Fig. 2B). In normal keratinocytes, exit from the basal layer is tightly coupled to terminal differentiation and the expression of cornified envelope proteins, such as IVL and TGM1. Consistent with this, when OKs were recovered from suspension and subjected to colony-forming assays, they lost their clonogenic potential. In contrast, although cancer cells expressed IVL and TGM1 in suspension the reduction in colony-forming ability after methylcellulose culture was less marked than that of OK (Fig. 2C and 2D).

**Figure 2.**
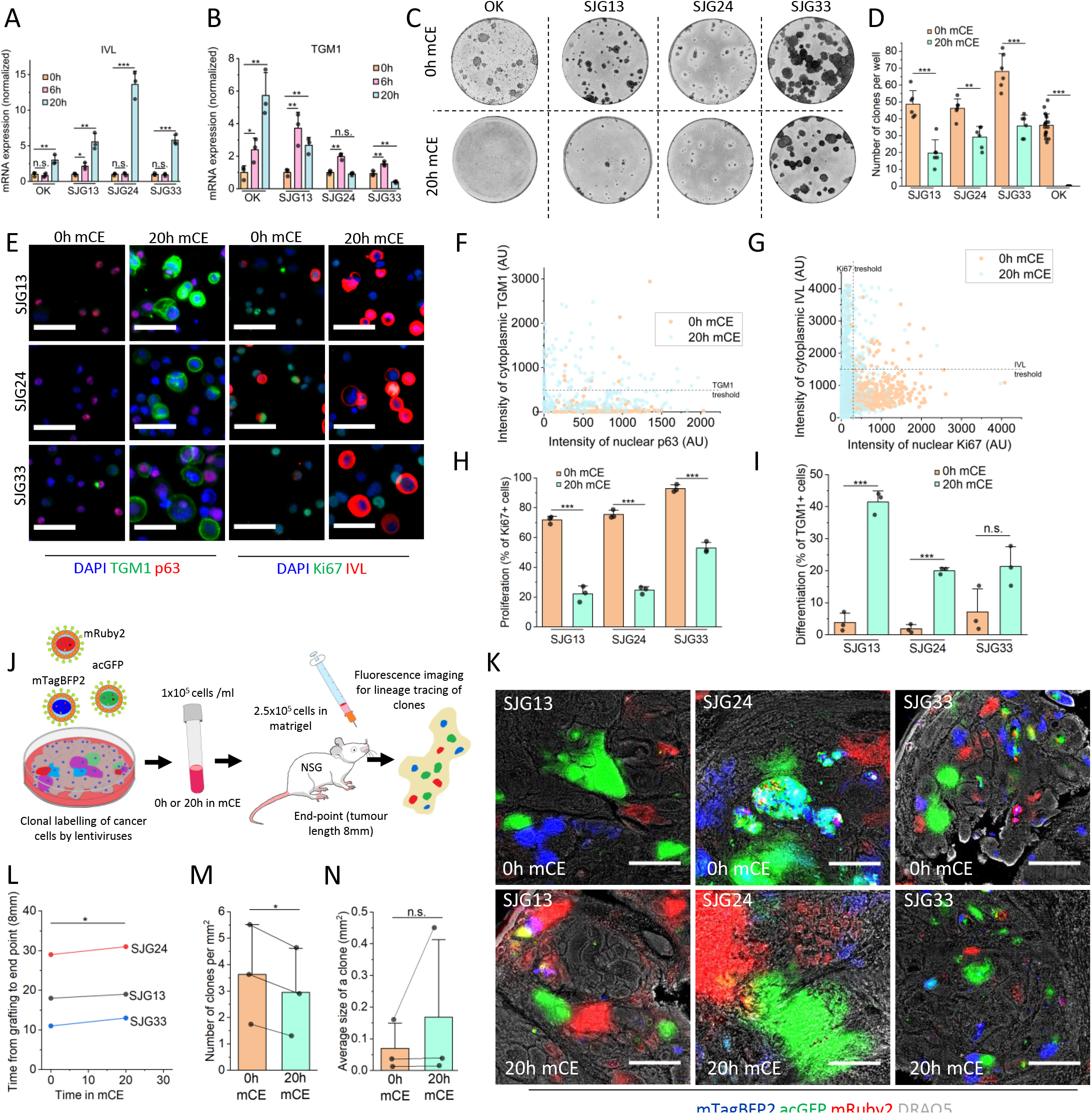
Uncoupling of differentiation and self-renewal in HNSCC cells. (A and B) qPCR analysis of IVL (A) and TGM1(B) expression in normal oral keratinocytes (OK) and SJG13, SJG24, and SJG33 cells kept in methylcellulose (mCE) for 0h, 6h, or 20h. Data represent mean ± SD of three biological replicates normalized to housekeeping genes. (C and D) Clonogenicity assay of OK, SJG13, SJG24, and SJG33 cells cultured in methylcellulose for 0h or 20h. Representative wells stained with Rhodanile Blue. (C) Data represent mean ± SD from 6 replicates (SJGs) and 18 replicates (OK) (D). (E) SJG13, SJG24, and SJG33 cells cultured in methylcellulose for 0h or 20h, disaggregated into single cell suspensions and stained for TGM1 (green) and p63 (red), Ki67 (green), or IVL (red), nuclei counterstained with DAPI (blue). Scale bars: 50 μm. (F and G) Quantification of TGM1 and p63 (F) and IVL and Ki67 (G) expression in disaggregated SJG13 cells after 0 h or 20 h in methylcellulose. Each dot represents one cell from a single experiment. (H) Percentage of proliferative (Ki67^+^) SJG13, SJG24 and SJG33 cells after 0h or 20h in methylcellulose. Mean ± SD from three independent SJG lines (; n = 3). (I) Percentage of differentiated (TGM1^+^) SJG13, SJG24 and SJG33 cells after 0h or 20h in methylcellulose. Mean ± SD from three biological replicates (n=3). (J) Schematic of the clonal lineage-tracing strategy. (K) Fluorescence imaging of clones in 40 μm thick cryosections of tumours derived from SJG cells cultured for 0h or 20h in methylcellulose. Nuclei were counterstained with DRAQ5 (white). Scale bars: 500 μm. (L) Time for tumours to reach the 8 mm endpoint in mice after methyl cellulose treatment. Mean ± SD from three independent SJG lines (SJG13, SJG24, SJG33; n=3 for 0h and 20h mCE). (M) Average number of clones per mm^2^ of tumour area at the end-point after 20h in methylcellulose (n=3), normalized to control (0 h; n=3). Mean ± SD from three independent SJG lines (SJG13, SJG24, SJG33). (N) Average clone area after 20h in methylcellulose (n=3), normalized to control (0h; n=3). Mean ± SD from three independent SJG lines (SJG13, SJG24, SJG33). Statistical significance was determined using two-tailed unpaired t-test (A, B, D, H, I) or two-tailed paired t-test (L, M, N). ***p < 0.0005; **p < 0.005; *p < 0.05.

To further characterize how the induction of differentiation affects the stem cell compartment of the cancer cell lines, we performed immunostaining of individual cells for differentiation markers (IVL and TMG1), the basal stem cell marker p63 and the proliferation marker Ki67. This analysis enabled simultaneous assessment of differentiation status and proliferative capacity within individual tumour cells (Fig. 2E-2I). We observe a major increase in Ki67 negative (Fig. 1H) and differentiated cells (Fig. 1I) after 24h in methylcellulose. However, a subset of cancer cells remained Ki67 positive or did not express differentiation markers (Fig. 2E-2I). Collectively, these findings indicate that although tumour cells partially respond to differentiation-inducing conditions, differentiation is not fully coupled to irreversible cell cycle exit (Parkinson 1985). Consequently, a subset of cells retains clonogenic potential despite expressing differentiation markers, suggesting the presence of differentiation-resistant tumour cell populations.

To determine the in vivo relevance of these findings, we established a lineage-tracing model based on fluorescent barcoding of single-cell clones by lentiviral expression of mRuby2, mTagBFP2, and acGFP. This approach enabled us to track the fate of clonogenic cancer cells in vivo and assess whether they were intrinsically resistant to differentiation cues. Typically, 30-40% of SJG24 cells and 10-20% of SJG13 as well as SJG33 cells expressed a fluorescent marker and a small number of cells were dual-labelled (Fig. 2K). There were no significant difference between the different fluorescent proteins and a small number of clones were dual-labelled. Following lentiviral labelling, cells were cultured in methylcellulose suspension for 24 hours and subsequently injected into mice (Fig. 2J). Tumours were allowed to grow to 8 mm in diameter before collection and analysis.

20h methylcellulose -treated cells showed a minor delay in tumour growth compared to cells that had not been placed in suspension (Fig. 2L). The clonal analysis revealed a minor decrease in clonal density (Fig. 2K and 2M) and increase in the average clone size, which was not statistically significant (2K and 2N). This strongly suggests that the HNSCC cells with high clonogenic capacity in vivo, and therefore responsible for tumour growth, are largely unaffected by transient detachment-induced differentiation signals.

### Afatinib promotes differentiation of the patient-derived HNSCC cells but fails to induce terminal differentiation of clonogenic tumour propagating cells

Having established that some HNSCC cells evade terminal differentiation, we next asked whether this is associated with resistance to pharmacological agents that promote differentiation. To address this, we performed a focused small-molecule screen on SJG13, SJG24, and SJG33 in vitro, based on pathways associated with keratinocyte differentiation commitment (Mishra et al., 2017). The panel included the histone deacetylase inhibitor valproic acid, the ErbB inhibitor afatinib, the ERK inhibitor VX-11e, the MEK inhibitor PD0325901, the PI3K inhibitor dactolisib, the mTOR inhibitor everolimus, and the IGFR inhibitor BMS-754807. Differentiation was assessed by induction of IVL mRNA expression (Fig. 3). Inhibitors were used at different concentrations depending their optimal active range (Fig. 3A-3C). Inhibition of the PI3K–mTOR axis by dactolisib or everolimus did not promote differentiation in any of the SJGs. In contrast, targeting the MAPK pathway induced differentiation efficiently. Both MEKi PD0325901 and ERKi VX-11e robustly increased differentiation marker expression in a concentration dependent manner (Fig. 3A-3C). ErbB inhibition by afatinib, which blocks upstream activation of both MEK and ERK signalling, also promoted differentiation as did IGFRi BMS-754807 albeit with lower efficacy. Valproic acid did not significantly induce differentiation in this model system (Fig. 3D). These results suggest that suppression of ErbB–MAPK signalling is the most effective strategy among those tested for inducing differentiation in these cells.

**Figure 3.**
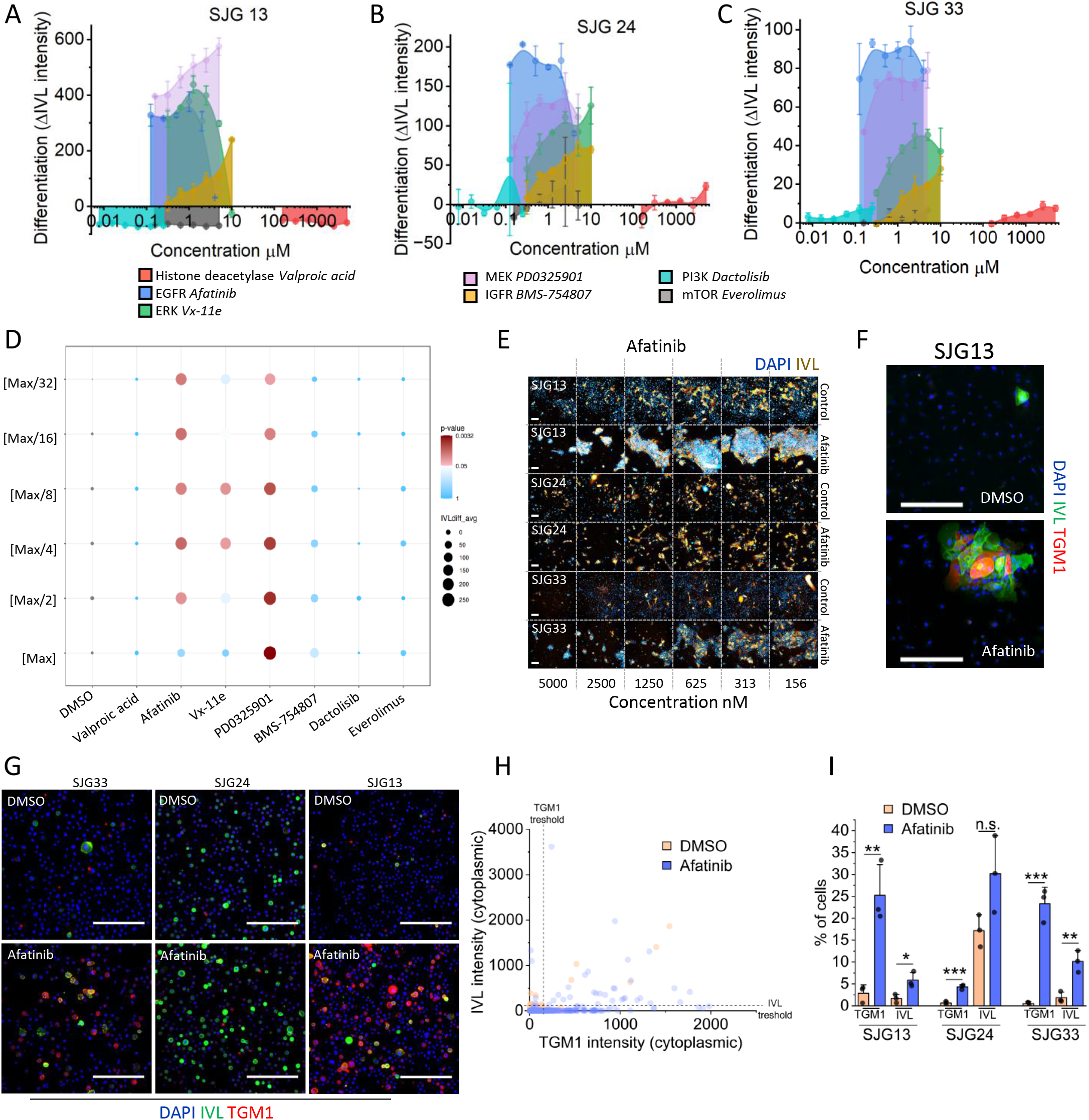
Inhibitors targeting the ErbB–MEK1/2–ERK1/2 pathway induce differentiation in HNSCC cells. (A–C) Quantification of IVL expression in SJG13 (A), SJG24 (B), and SJG33 (C) cells following treatment with different concentrations of small-molecule inhibitors for 48h. Cells were stained with an anti-IVL antibody and analysed using the Operetta high-throughput imaging system. Mean ± SD from three biological replicates is shown (n = 3). Y-axis shows the fluorescence difference (increase or decrease) compared to control. (D) Integrated statistics of Operetta analysis and ranking of the most effective drugs and concentrations across SJGs (SJG13, SJG24, and SJG33; n = 3). Y-axis is a relative drug concentration. Max concentrations of the drugs: [valproic acid] 5000µM, [afatinib] 4µM, [Vx-11e] 10µM, [PD0325901] 5µM, [BMS-754807] 10µM, [Dactolisib] 0.25µM, and [everolimus] 5µM. (E) Operetta fluorescence imaging of SJG13, SJG24 and SJG33 treated with different afatinib concentrations for 48h and stained for IVL (brown) with nuclei counterstained with DAPI (blue). Scale bars: 200 μm. (F) Confocal fluorescence imaging of SJG13 treated with 200nM afatinib for 48h and stained for IVL (green) and TGM1 (red) with nuclei counterstained with DAPI (blue). Scale bars: 250 μm. (G) SJG13, SJG24, and SJG33 cells treated with afatinib for 48h, disaggregated into single cell suspensions and stained for IVL (green) and TGM1 with nuclei counterstained with DAPI (blue). Scale bars: 200 μm. (H) Quantification of TGM1 and IVL expression by confocal in disaggregated SJG13 cells after treating cells with 200nM afatinib for 48h. Each dot represents one cell from a single experiment. (I) Percentage of differentiated (IVL?) cells after treating cells with 200nM afatinib for 48h. Mean ± SD of three biological replicates from disaggregated SJG13, SJG24, and SJG33 cells (n = 3). Analysis was done by confocal imaging of disaggregated cells. Statistical significance was determined using two-tailed unpaired t-test. ***p < 0.0005; **p < 0.005; *p < 0.05.

We selected ErbBi afatinib for further investigation given its established clinical use in HNSCC. Although afatinib has not demonstrated a significant improvement in overall survival in patients, it has shown clinical activity by prolonging progression-free survival (Guo et al., 2019). The clinical trials with afatinib have shown that its plasma concentration level in humans remains under 200nM when using the standard treatment (Wind et al., 2017). Treatment of the SJG lines with 200nM afatinib lead to increased expression of TGM1 and IVL (Fig. 3F-3I). Nevertheless, a substantial fraction of cells did not upregulate differentiation markers following afatinib treatment (Fig. 3G-3I), suggesting the presence of a differentiation-resistant subpopulation, as was the case in the methylcellulose suspension assay.

To determine whether ErbB inhibition targets the cells that are tumorigenic in vivo, we employed random lentivirus-based labelling strategy as in the case of methyl cellulose assay (Fig. 2J and 4A). SJG 13, SJG24, SJG33 cells were transfected with lentivirus and subsequently treated with 200 nM afatinib for 48h. Following treatment, the cells were injected into mice to assess tumour formation. No significant difference in tumour size was observed between control and afatinib-treated groups (Fig. 4B-4C). ErbB inhibition did not alter either clone area or clone size between treated and control groups (Fig. 4E-4F). These findings demonstrate that ErbB inhibition is insufficient to drive irreversible cell cycle withdrawal and terminal differentiation in tumorigenic cells with in vivo tumour-forming capacity.

**Figure 4.**
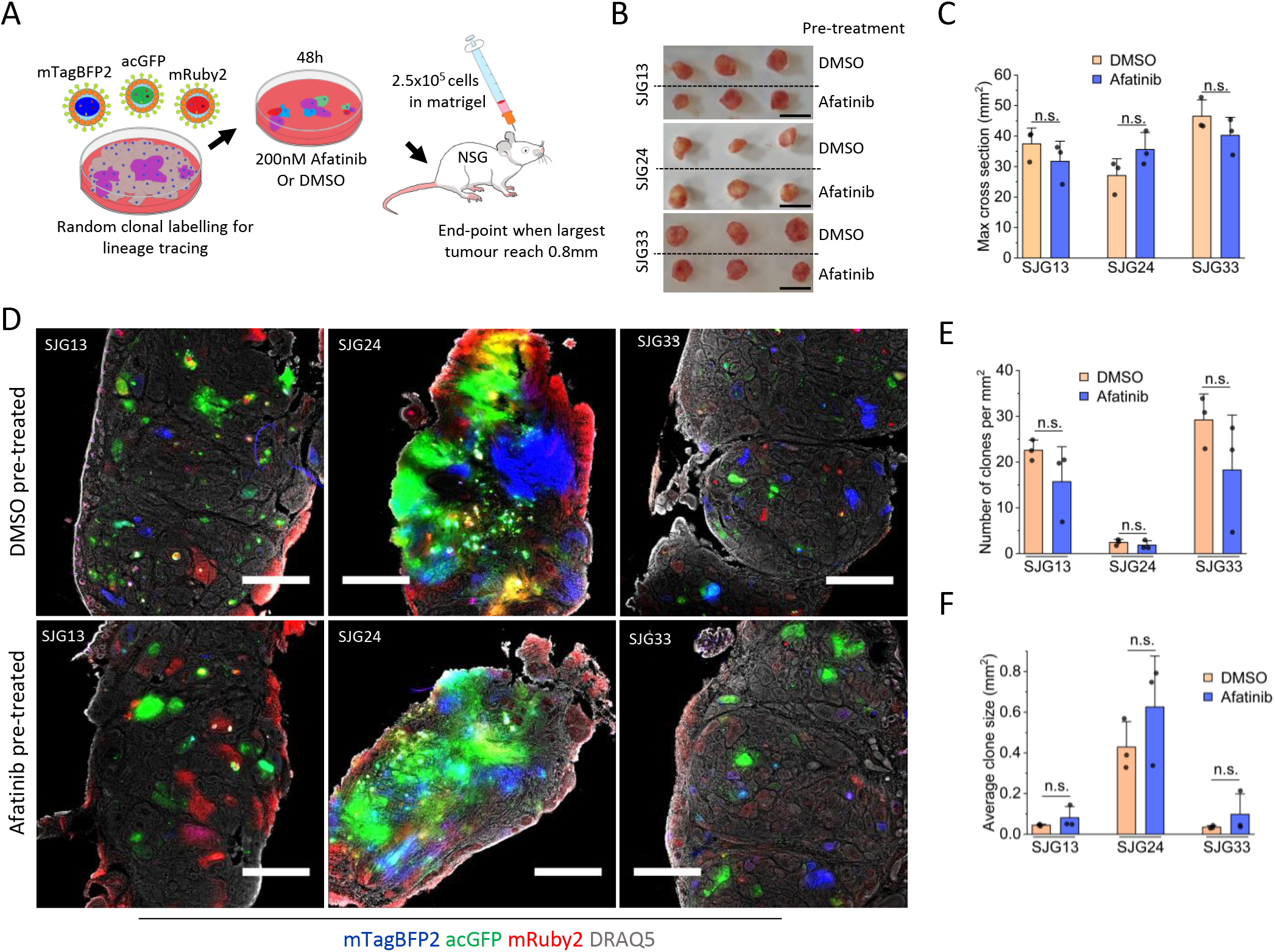
Induction of differentiation leads to a gradual reduction in the tumorigenic potential of HNSCC cells. (A) Schematic of the clonal lineage-tracing experiment using visual barcoding after afatinib pre-treatment. (B) Macroscopic images of mouse cheek xenograft tumours harvested from control (DMSO) or afatinib pre-treated SJG13, SJG24, and SJG33. Scale bars: 10mm. (C) Quantification of xenograft tumour sizes derived from control (DMSO; n=3) or afatinib pre-treated (200nM, 48h; n=3) SJG13, SJG24, and SJG33. (D) Fluorescence imaging of clones in 50 μm thick cryosections of tumours derived from SJG cells (SJG13, SJG24, and SJG33) pre-treated with DMSO or afatinib (200nM, 48h); DRAQ5 nuclear counterstain (white). Scale bars: 1000μm. (E) Number of clones per tumour in DMSO (n=3) or afatinib pre-treated tumours (200nM, 48h; n=3). Mean ± SD is shown. (F) Average clone area in DMSO (n=3) or afatinib pre-treated tumours (200nM, 48h; n=3). Mean ± SD is shown. Statistical significance was determined using two-tailed unpaired t-test. ***p < 0.0005; **p < 0.005; *p < 0.05.

### Plasticity of squamous differentiation drives resistance to afatinib mediated differentiation

These observations prompted us to investigate whether afatinib treatment alters the tumorigenic capacity of cells depending on their differentiation status. To directly assess this, we transfected SJG13 cells with an IVL promoter–driven fluorescent reporter (IVLmCherry; Hiratsuka et al., 2020) as a differentiation reporter, together with a constitutive GFP marker serving as a global lineage tracer (Fig. 5). This dual-labelling strategy enabled identification and prospective isolation of cells expressing IVL at the time of fluorescence-activated cell sorting (FACS), while GFP expression allowed tracking of the total progeny that afatinib-treated cells formed in vivo.

**Figure 5.**
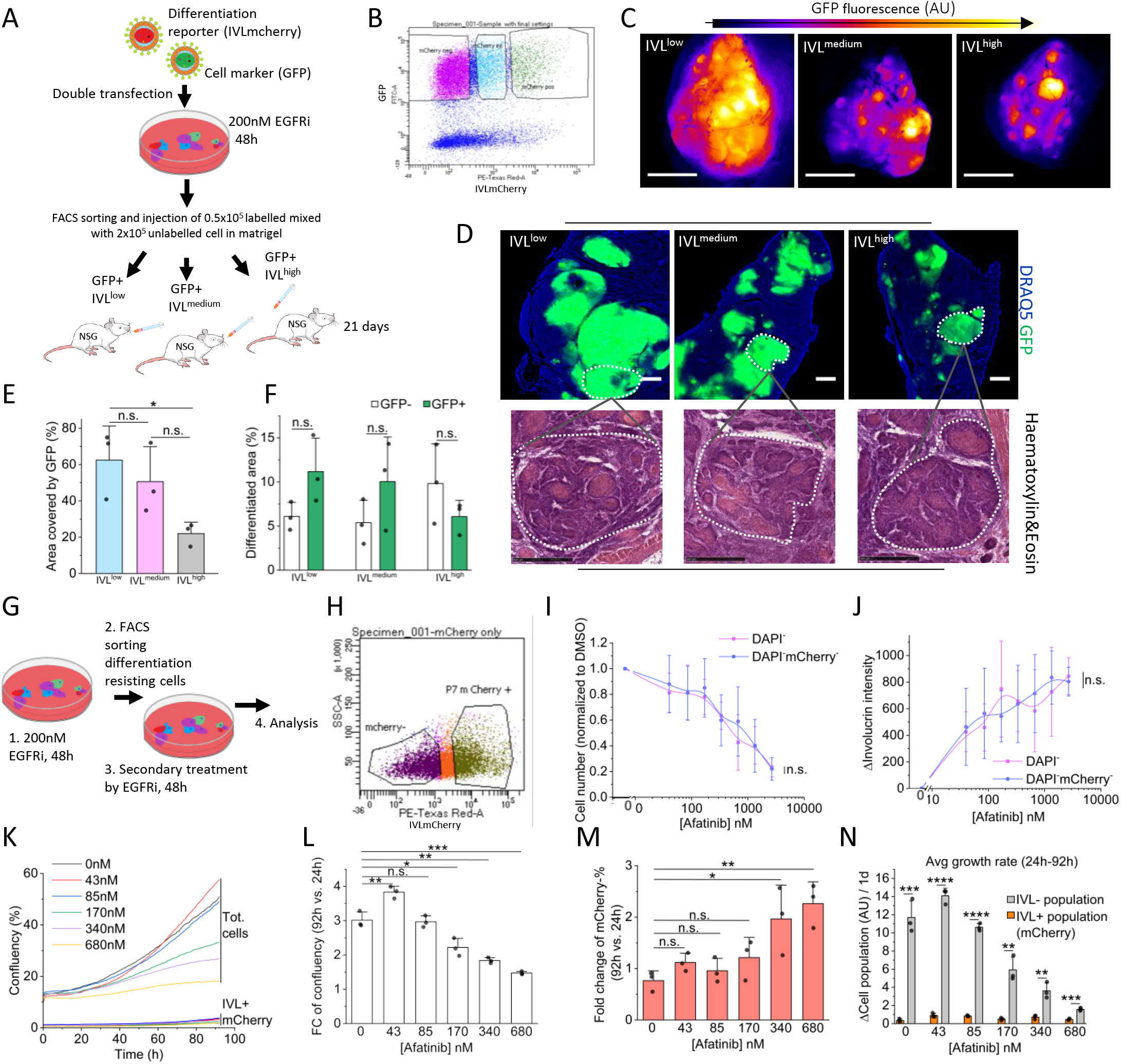
Afatinib fails to induce terminal differentiation in clonogenic tumour-initiating cells. (A) Schematic of the lineage-tracing experiment using IVLmCherry reporter. (B) FACS strategy to sort IVL^low^, IVL^medium^, and IVL^high^ cell fractions after 48h afatinib pre-treatment of SJG13 cells. (C) Whole mount fluorescent (GFP) images of intact mouse cheek tumours from IVL^low^, IVL^medium^, and IVL^high^ cell fractions. (D) Confocal (GFP green, nuclear DAPI stain blue) and H&E images of 50 μm thick cryosections from the tumours from IVL^low^, IVL^medium^, and IVL^high^ cell fractions. Scale bars: 500 μm (E) Quantification of tumour area covered by IVL^low^, IVL^medium^, and IVL^high^ GFP positive cells. Mean of each group (n=3) ± SD is shown. (F) Quantification of differentiating tumour areas derived from GFP positive (IVL-fractionated) or GFP negative (non-fractionated) cells. Mean of each group (n=3) ± SD is shown. (G) Schematic of the serial testing of afatinib induced differentiation using IVLmCherry reporter in SJG13 cells. (H) FACS strategy to sort IVL^low^ and total cell fraction after 48h afatinib treatment of SJG13 cells by using IVLmCherry reporter. (I and J) Cell number (I) and IVL expression by antibody staining (J) after treating differentiation-resistant viable cells (DAPI negative, IVLmCherry) and total viable cells (DAPI negative) with different afatinib concentrations for 48h. control. Mean of independent biological replicates (n=3) ± SD is shown. (K) Area covered by IVLmCherry positive or SJG13 negative cells treated with afatinib as a function of time. Mean of independent biological replicates (n=3) is shown. (L, M, N) Change of total confluence (L) and differentiation area (IVLmCherry positive; M) as well as average growth rate of IVLmCherry positive and negative populations (N) between attachment (one day after plating) and 92h after plating in the presence of different afatinib concentrations. Mean of independent biological replicates (n=3) ± SD is shown. Statistical significance was determined using two-tailed unpaired t-test. ***p < 0.0005; **p < 0.005; *p < 0.05.

Cells were treated with 200 nM afatinib for 48h. Following the treatment, cells were sorted based on IVLmCherry intensity into three fractions: high, intermediate, and low/negative expressers (Fig. 5A-5B). All the sorted cells were viable based on DAPI negativity. When sorted cells were mixed with non-sorted GFP-negative cells and transplanted, there was a clear inverse relationship between IVL expression and tumorigenic capacity (Fig. 5C-5E). Cells with high IVL expression formed markedly smaller clones compared to IVL-low cells in vivo. Furthermore, the data indicate that differentiation and reduction of tumorigenic potential are gradual rather than binary, since a subset of IVL high cells retained the capacity to generate substantial progeny in vivo (Fig. 5C-5E).

Histological analysis of tumour sections by H&E after imaging the fluorescence revealed that tumours displayed a similar overall degree of differentiation in vivo regardless of the level of IVLmCherry expression in the initial sorted population (Fig. 5D and Fig. 5F). To further investigate this phenomenon, we examined the behaviour of sorted populations in vitro (Fig. 5G-5J). After an initial treatment with afatinib, IVL-negative/low cells were sorted and re-tested for their sensitivity to afatinib in terms of growth and differentiation, compared to the non-sorted population (Fig. 5I-5J). We observed no significant difference in sensitivity to afatinib between the IVL-negative/low subpopulation and the total cell population (Fig. 5I). Moreover, the cells in the IVL low/negative fraction retained the ability to give rise to progeny that expressed IVL after treatment (Fig. 5J). These findings indicate that although some cells resist afatinib-mediated induction of differentiation, they retain the ability to differentiate and retain sensitivity to inhibition of ErbB. Importantly, even at supra-clinical concentrations, an afatinib resistant subpopulation persists, highlighting the intrinsic resistance to ErbB-induced differentiation (Fig. 5K-5N and Fig. 3A). Together, these results suggest that afatinib has a limited capacity to enforce terminal differentiation of HNSCC cells.

## Discussion

Normal epithelial keratinocytes undergo differentiation after detachment from the basement membrane, and terminal differentiation is coupled to loss of self-renewal capacity. The interaction between epithelial stem cells and their niche regulates differentiation dynamics during development and under homeostatic conditions (Sipilä et al., 2022; Zijl et al., 2022; Coulombe and Wickström, 2021). In this work, we demonstrate that patient-derived cancer cells cultured on J2-3T3 cells using an epithelial stem cell culture method respond to differentiation induction in a highly heterogeneous manner: some cells differentiate but others remain highly clonogenic. In xenografts, individual tumour cell lines recapitulate the epithelial organization of the primary tumours.

Our findings using the differentiation reporter IVLmCherry further support the concept that differentiation commitment in cancer cells is not governed by a strict binary on/off switch, but that cells may occupy a range of intermediate states along a spectrum. The most tumorigenic cells in vivo were enriched within the IVLmCherry–low fraction, whereas cells with high IVLmCherry expression exhibited markedly reduced tumorigenic capacity in vivo. Nevertheless, some IVL-GFP–positive cells retained the ability to generate progeny in vivo, indicating that expression of differentiation markers does not necessarily equate to irreversible loss of tumour-initiating potential. Our experiments do not rule out the possibility that some of the tumour cells in the IVL-high fraction de-differentiated back to a more stem cell–like state in tumours in vivo (Donati et al., 2017). However, it is possible that ErbBi by afatinib treatment changed only the expression of differentiation markers in a reversible manner. Importantly, our approach, to use random cell labelling, measures the functional consequence of the drug treatment to in vivo clonogenic capacity independent on marker expression.

Our findings reveal important conceptual similarities between APL and HNSCC. In APL, differentiation therapy using ATRA induces leukemic cell differentiation and can lead to durable remissions. However, a full curative effect depends on effectively targeting the subpopulation of cells capable of resisting or escaping differentiation. Similarly, in our patient-derived HNSCC models, pharmacological inhibition of ErbB causes a subpopulation to differentiate. A distinct subset of cells fails to undergo stable differentiation, and our data demonstrate that these differentiation-resistant cells correspond to the most highly clonogenic and tumorigenic population in vivo. While conventional chemotherapy may target rapidly proliferating cells, it remains unclear whether the cells with high clonogenic and self-renewing capacity proliferate at sufficient rates to be effectively eliminated by such treatments (Chen et al., 2017). Thus, achieving curative outcomes in HNSCC may require a deeper understanding of the mechanisms by which this subpopulation blocks or resists terminal differentiation. Importantly, our work provides a conceptual platform to understand the differentiation dynamics of HNSCCs and to develop therapeutic strategies that overcome the differentiation plasticity-driven resistance either by forcing cells to undergo irreversible differentiation or by directly eliminating these cells.

## Materials and Methods

### Animal procedures

All animal work was carried out under a UK Government Home Office licence (PP70/8474 or PP0313918) and was approved locally by the Animal Welfare and Ethical Review Body of King’s College London (UK). The mouse line used for the tumour xenografting studies was NOD.Cg-Prkdc^scid Il2rg^tm1Wjl/SzJ (NSG®), obtained from Charles River (RRID:IMSR_JAX:005557). Only adult female mice were used. For tumour grafting, cells were suspended in ice-cold Matrigel and injected into the mouse cheek via the upper lip using 29–30G insulin syringes. A total of 30 µl of Matrigel containing 0.2–0.5 million cells was injected. The procedure was performed under isoflurane anaesthesia, and mice received a subcutaneous injection of buprenorphine as an analgesic (0.15 mg/kg). Tumour growth and mouse weights were monitored 2–3 times per week, and the endpoint was reached when a tumour (or the largest tumour in the cohort) reached a size of 8 mm. For tissue harvesting, animals were sacrificed by cervical dislocation or by exposure to increasing concentrations of CO□. Tumours were imaged and preserved in OCT (optimal cutting temperature compound) at ™80 °C.

### SJG cell cultures and colony forming assay

SJG cells were isolated and cultured as described previously (Broad et al. 2026; Hayes et al. 2016; Goldie et al. 2012). Briefly, 0.25 million SJG cells or normal oral keratinocytes (OK) were seeded on top of a mitotically inactivated (mitomycin C) J2-3T3 feeder layer (1.8million cells) in T75 flasks. Flasks were typically split when reaching 80–90% confluence by detaching the cells with 0.25% trypsin/EDTA. SJGs and OKs were cultured in complete FAD medium (3:1 ratio Dulbecco’s Modified Eagle’s Medium (DMEM) and Ham’s F-12 medium, 5mM□L-glutamine, 0.18mM adenine, 10% FBS, 0.5μg/ml hydrocortisone, 5μg/ml insulin, 0.1nM cholera toxin, EGF 10ng/ml, 100□IU/ml penicillin, and 100μg/ml streptomycin) in a cell culture incubator (5% CO□, 37°C) and the medium was changed three times per week. J2-3T3 cells were expanded in DMEM (glucose 4.5g/l) supplemented with 10% bovine serum, 5mM□l-glutamine, 100□IU/ml penicillin, and 100μg/ml streptomycin. Passages under 15 were used for all cells. For the colony-forming assay, 250 SJG cells or 1000 OK cells were seeded into each well of a 6-well plate covered with a mitomycin-treated J2-3T3 feeder layer. Media were changed three times per week, and the assay was terminated 10–12 days after seeding. Feeders were removed by washing with PBS. SJG and OK colonies were fixed with 4% paraformaldehyde (PFA, 10min) and stained with 1% Rhodanile Blue (1:1 mixture of Rhodamine B and Nile Blue chloride). The number of colonies was counted manually.

### Lentiviral transfections

Lentiviral particles were produced by using HEK-293 cells and the second generation packaging system as described earlier (Oulès et al., 2020). The following plasmids were used: pLenti-Involucrin-mCherry (Hiratsuka et al., 2020), pLNT-SFFV-mRuby2 (a gift from Violaine Sée; Addgene plasmid#87217; http://n2t.net/addgene:87217;RRID:Addgene_87217), mTAGBFP2 WC visual barcode (a gift from Ravid Straussman; Addgene plasmid#158666; http://n2t.net/addgene:158666; RRID:Addgene_158666), acGFP WC visual barcode (a gift from Ravid Straussman; Addgene plasmid#158672; http://n2t.net/addgene:158672; RRID:Addgene_158672), pNano-Lenti-bGHpolyA-SFFV-UbiC-EGFP (OXGENE). SJG cells were transduced in complete FAD containing 5μg/ml polybrene (EMD Millipore) and the medium was changed in 16h. SJG13, transduced by pLenti-Involucrin-mCherry, were selected and maintained in FAD supplemented with 2 µg/mL puromycin.

### Drug treatment

Cells were plated one day before drug treatments in 96-well plates or other tissue culture vessels in complete FAD medium at a density of 3125 cells/cm^2^. To start the treatment, the medium was replaced with drugs diluted in FAD. After 48h, cells were either fixed with 4% PFA or trypsinized for poly-L-lysine slides and for injection into mice. The stock solutions of the inhibitors were diluted in DMSO and following inhibitors were used: valproic acid (Merck), afatinib (APExBIO), Vx-11e (Selleckchem), PD0325901 (Selleckchem), BMS-754807 (TargetMol Chemicals), Dactolisib (Selleckchem), and everolimus (APExBIO). Drug concentrations are specified in the figures. Afatinib concentration was 200nM unless indicated otherwise. In some experiment, the expansion of cells plated on 6-well plates with different afatinib concentrations was measured by Incucyte (phase contrast and mCherry fluorescence; Sartorius). SJG13-IVLmCherry and SJG13-IVLmCherry-GFP cells were detached after 48h afatinib treatment and sorted by FACSAria III Cell Sorter (BD Biosciences) using DAPI negativity as a marker of viable cells.

### Methyl cellulose treatment

The methylcellulose assay was performed as described earlier (Adams and Watt, 1989). Briefly, detached SJG cells or oral keratinocytes were suspended (0.1 million cells per ml) in complete FAD medium containing 1.45% dissolved methylcellulose (viscosity 4000 cP). Cell suspensions were placed into polypropylene round-bottom bacterial test tubes with semi-open lids (Corning) to prevent cells from attaching to the tube and to maximize gas exchange. Cell suspensions were kept in a cell culture incubator (5% CO□, 37 °C) at different time points. Cells were harvested from methylcellulose by diluting the suspension with PBS (1:10) followed by centrifugation.

### qPCR

RNA was isolated from the samples using the RNeasy Mini Kit (Qiagen, 74104), and cDNA was synthesized using the QuantiTect Reverse Transcription Kit (Qiagen, 205311) according to the instructions provided by the manufacturer. qPCR was performed using Fast SYBR® Green amplification (Applied Biosystems, 4385612). The following primers were used (Merck/Sigma): Hs_IVL_F (GCCTCAGCCTTACTGTGAGT), Hs_IVL_R (TGTTTCATTTGCTCCTGATGG), Hs_TGM1_F (GCACCACACAGACGAGTATGA), Hs_TGM1_R (GGTGATGCGATCAGAGGATTC), Hs_ATP5B_F (AGGCTGGTTCAGAGGTGTCT), Hs_ATP5B_R (TGGGCAAACGTAGTAGCAGG), Hs_TBP_F (GTGACCCAGCATCACTGTTTC), Hs_TBP_R (GAGCATCTCCAGCACACTCT), Hs_RPL13A_F (AACAGCTCATGAGGCTACGG), Hs_RPL13A_R AACAATGGAGGAAGGGCAGG. IVL and TGM1 expression levels were normalized to the housekeeping genes (ATP5B, RPL13A, and TBP).

### Disaggregation of the cells onto poly-L-lysine slides

After cells were trypsinized or harvested from methylcellulose, they were washed and suspended in complete FAD medium (5 million cells per ml). 10–50 µl of the suspension was pipetted onto poly-L-lysine slides (Thermo Scientific). After allowing the cells to attach for 20 min in a cell culture incubator (5% CO□, 37°C), the medium was aspirated, the cells were fixed with 4% PFA (10min, RT), and processed for immunofluorescence staining.

### Histological and immunofluorescence staining

Cryopreserved tumour samples were sectioned into 12 µm or 40 µm slices using a cryostat. Haematoxylin and eosin (H&E) staining was performed, and the pathological features of the samples were assessed by a trained expert oral pathologist under bright-field microscopy. Cryosections, as well as cells attached to culture vessels or poly-L-lysine slides, were fixed with 4% PFA (10 min, RT), followed by permeabilization with 0.5% Triton X-100 in PBS (10 min, RT). After permeabilization, samples were blocked with blocking buffer (10% FBS, 0.25% fish gelatin, and 3% BSA) and incubated with primary antibody overnight at 4 °C. Primary antibodies were diluted in blocking buffer, and the following antibodies and dilutions were used: anti-Ki67 (1:500-1:1000, SP6, Abcam), anti-IVL (1:1000 clone SY7 or DH1B6), anti-TGM1 (1:500 clone BC1), anti-K14 (1:500, Poly19053, BioLegend), and anti-p63-alpha (1:500, D2K8X, Cell Signaling Technology). Samples were then treated with secondary antibodies (Alexa 488, Alexa 555, or Alexa 647; Invitrogen). Nuclei were counterstained with DAPI or DRAQ5 in PBS (10 min, 5 μM in PBS; Abcam). Cells were washed three times for 5 min with PBS after antibody incubation and nuclear staining steps. Finally, 12 µm sections were mounted in ProLong™ Gold Antifade Mountant (Thermo Fischer Scientific), 40 µm tumour sections were mounted in glycerol, and attached cells were kept in PBS before imaging. Some 40 µm tumour sections were imaged after fixing and DRAQ staining, and H&E was performed after imaging. Samples were imaged using Nikon A1 upright scanning confocal microscope, Perkin-Elmer Operetta CLS High-Content Imaging System Operetta, or Hamamatsu NanoZoomer slide scanner.

### Image analysis

Fluorescence intensity, clonal analysis, and cell counting were performed using Operetta Harmony™ software, Fiji (ImageJ), and QuPath. Thresholds for positive and negative cells were manually determined. Clone areas were manually defined. The area of tumour cross-sections were quantified from tumour photographs with measurement scales using Fiji (ImageJ).

## Data representation and statistical analysis

All graphs and statistical analyses were generated using R or OriginLab. Statistical significance was determined using a two-tailed Student’s t-test (paired or independent means) from three biological replicates, unless otherwise indicated. Schematics were created using Microsoft Paint, and figures were assembled using Microsoft PowerPoint. ChatGPT (OpenAI) was used to correct grammar and improve the readability of the text.

## Acknowledgements

We are thankful for the financial support from Cancer Research UK (C219/A23522; F.M.W.), the Medical Research Council (G1100073; F.M.W.), and the Wellcome Trust (096540/Z/11/Z; F.M.W.), Department of Health via the National Institute for Health Research comprehensive Biomedical Research Centre (BRC) award to Guy’s & St Thomas’ National Health Service Foundation Trust in partnership with King’s College London and King’s College Hospital NHS Foundation Trust, and from the Finnish Cultural Foundation for supporting the postdoctoral fellowship of K.S. We would also like to thank Prof. Matthew Garnett (Sanger Institute) for the initial drug screening and Dr. Toru Hiratsuka for pLenti-Involucrin-mCherry.

